# Chemotherapeutic cardiotoxicity is associated with elevated β1-adrenergic receptor density

**DOI:** 10.1101/2020.09.17.301689

**Authors:** Manveen K Gupta, Elizabeth E. Martelli, Kate T. Stenson, Sathyamangla V. Naga Prasad

**Author notes:** **Address correspondence to:** Sathyamangla V. Naga Prasad, PhD, FAHA or Manveen K. Gupta, PhD MBA, or, Department of Cardiovascular Medicine, Lerner Research Institute, Cleveland Clinic, 9500 Euclid Avenue, Cleveland, OH, 44195. No industry relationship to disclose.

## Abstract

**Objective:** To understand the underlying pathways that promote cardiotoxicity following chemotherapy.

**Background:** Anthracyclines are associated with cardiotoxicity which could be potentiated with use of complementary agents (like anti-ERBB2 inhibitors) which together afford robust anti-neoplastic effects. Anthracyclines lead to oxidative stress and thought to induce cardiotoxicity. However, interventions reducing oxidative stress in patients have been unsuccessful suggesting mechanisms beyond oxidative stress. Despite β-adrenergic receptors (βARs) being key regulators of cardiac function, nothing is known about their role in chemotherapy-mediated cardiotoxicity.

**Methods:** β1 and/or β2-AR density was assessed in end-stage human heart failure patient samples either due to anthracycline cardiotoxicity or non-anthracycline dilated cardiomyopathy (DCM). Since ERBB2 inhibition is integral to overall chemotherapeutic arsenal, we assessed β1- and/or β2-AR density, cardiac function by echocardiography and immunohistochemistry in mice following ERBB2-specific inhibitor AG825.

**Results:** Selective increase in cardiac β1AR density is observed in end-stage human heart failure patient samples due to anthracycline cardiotoxicity as well as in ERBB2 inhibitor-treated mice.

**Conclusions:** Elevated β1AR density may be the key common underlying mechanism which is altered in response to chemotherapy promoting cardiac dilation of otherwise healthy hearts.

**Highlights:** In contrast to downregulation of β1-adrenergic receptors (β1AR) in end-stage human heart failure, anthracycline cardiotoxicity-mediated failure is associated with selective increase in β1AR density.

ERBB2 inhibitor (AG825) treatment in mice results in cardiac dilation and selective rise in β1AR density showing that increased β1AR density in the heart could be a common mechanism underlying cardiotoxicity.

## Introduction

Anthracyclines have been the main stay therapy for cancer but, is associated with significant cardiotoxicity^1, 2^. It is estimated that ~9% of the patients on anthracyclines will display cardiotoxicity ranging from dilated cardiomyopathy (DCM), congestive heart failure, atrial fibrillation to subclinical structural changes reflecting patient susceptibility^1^. Retrospective studies have shown that majority of the patients develop cardiac dysfunction/failure spanning the New York Heart Association (NYHA) Classes I through IV heart failure classification^3^. Given that conventional anthracyclines causes cardiotoxicity, next generation of targeted cancer therapeutics were generated that had specific pharmacologic action like trastuzumab (humanized antibody directed against HER2/ERBB2 (Epidermal growth factor receptor 2/Erythroblastic oncogene B, ERBB2), sunitinib (Vascular endothelial growth factor (VEGF) inhibitor), dasatinib (Break point cluster (Bcr) chimera with Abl tyrosine kinase (BCR-ABL) inhibitor) to name a few^1^. However, majority of these targeted therapies are still associated with cardiotoxicity^1^. Despite the knowledge that chemotherapy increases oxidative stress resulting in lipid peroxidation and mitochondrial dysfunction, use of anti-oxidants in patients was not cardio-protective^4^. This shows that additional mechanisms underlie chemotherapy-induced cardiotoxicity.

β-adrenergic receptors (βARs) are powerful regulators of cardiac function, but less is known about their roles in cardiac remodeling following cardiotoxicity. βAR family is subdivided into β1AR, β2AR and β3AR^5^ based on their pharmacological properties with β1AR being the major regulator of cardiac function^5^. Consistently, end-stage human heart failure patient samples are characterized by selective downregulation of β1ARs associated with DCM and heart failure^6^. Given the key role of β1ARs in cardiac function, we assessed whether alterations in β1AR density underlies cardiotoxicity with anthracycline (using end-stage anthracycline-mediated human heart failure patient samples) or ERBB2 targeting (by treating mice with ERBB2-specific inhibitor (AG825)) representing a common mechanism for deleterious cardiac remodeling.

## Methods

### Human heart failure samples

Procurement of the end-stage human heart failure samples were approved by the institutional IRB. All the patient samples (non-failing, DCM or DCM due Adriamycin cardiotoxicity) were de-identified. Gender, age and clinical characteristics are shown in **Table 1**.

**Table 1:** Patient Characteristics

|  | Age | Gender | Race | Cause of death | Diagnosis | % LVEF | Medications prior to the event | Acutely administered medications |
| --- | --- | --- | --- | --- | --- | --- | --- | --- |
| NON-FAILING |  |  |  |  |  |  |  |  |
| 1 | 57 | F | W | CVA | - | ND | - | DOP |
| 2 | 61 | F | W | CVA | - | 50 | - | DOP, NE |
| 3 | 60 | F | W | Sub-AB | - | 65 | - | DOB, NE, PHE, THY |
| 4 | 62 | F | W | CVA | - | ND | - | NE, PHE, VP |
| 5 | 64 | F | W | CVA | - | 70 | - | NE, THY, VP |
| 6 | 62 | F | B | CVA | - | 67 | - | DOP, NE, THY, VP |
| DILATED CARDIOMYOPATHY (DCM) HEART FAILURE |  |  |  |  |  |  |  |  |
| 1 | 60 | F | W | - | DCM-Viral | 10 | AM, DIG, DOB | - |
| 2 | 61 | F | W | - | DCM | 10 | CARV, DIG, RAMP | - |
| 3 | 63 | F | B | - | DCM | 18 | CARV, DIG | - |
| 4 | 62 | F | W | - | DCM | 14 | CARV, | - |
| 5 | 63 | F | W | - | DCM | 31 | CARV, QUIN | - |
| 6 | 58 | F | B | - | DCM | 18 | AM, DIG, MIL | - |
| CHEMOTHERAPEUTIC (ADRIAMYCIN) HEART FAILURE |  |  |  |  |  |  |  |  |
| 1 | 62 | F | W | - | DCM-Chemo | 12 | CAPT, DIG, MIL | - |
| 2 | 63 | F | B | - | DCM-Chemo | 10 | CARV, LOS | - |
| 3 | 63 | F | W | - | DCM-Chemo | 8 | DIG, LIS, MIL | - |
| 4 | 58 | F | W | - | DCM-Chemo | 19 | - | - |
| 5 | 61 | F | W | - | DCM-Chemo | ND | AM | - |

ABBREVIATIONS: AM- Amiodarone, B- Black, CAPT- Captopril, CARV- Carvedilol, CVA- Cerebral vascular accident, DCM- Dilated cardiomyopathy, DOB- Dobutamine, DOP- Dopamine, F- Female, LIS- Lisinopril, LOS- Losartan, LVEF- Left ventricular ejection fraction, MIL- Milrinone, ND- Not determined, NE- Norepinephrine, PHE- Phenylephrine, QUIN- Quinapril, RAMP- Ramapril, Sub-AB- Sub-Arachnoid Bleed, THY- Levothyroxine, VP- Vasopressin, W- White.

### Plasma membrane isolation and βAR density

Plasma membranes were isolated as described previously ^7^. Pellet representing the plasma membrane fraction was resuspended in 75 mM Tris-HCl pH 7.5, 2 mM EDTA, and 12.5 mM MgCl_2_ for assess βAR density. βAR density was determined by incubating 20 μg of the membranes from mouse or end-stage human heart failure samples with saturating concentrations of ^125^I-Cyanopindolol. 100 μM propranolol was used for assessing non-specific binding and β2AR specific antagonist ICI 181,551 was used for determining β2AR density. These were subtracted from the non-specific propranolol values to determine β1AR density.

### Experimental Animals

Male and female C57BL/6 mice from Jackson labs were used for the studies. Animals were handled according to the approved protocols and animal welfare regulation of IACUC at Cleveland Clinic following the approved NIH guidelines.

### ERBB2 inhibitor studies

ERBB2-specific inhibitor AG 825 dissolved in DMSO and diluted in saline was used at a final concentration of 1 mg/kg/day through min-osmotic pump (ALZET model 2002) for 14 days.

### Echocardiography

Echocardiography was performed on anesthetized mice using a VEVO 770/VEVO 2100 (VISUALSONICS) echocardiographic machine as previously described^8^. M-mode recording was used to obtain functional parameters.

### Statistics

Data are expressed as mean ± SEM. Statistical comparisons were performed using an unpaired Student’s *t*-test for two samples comparison (like binding studies) and analysis of variance (ANOVA) was carried out for multiple comparisons (like paired echocardiography analysis). Post-hoc analysis was performed with a Scheffe’s test. For analysis, a value of * p< 0.05 was considered significant.

## RESULTS AND DISCUSSION

### Adriamycin-mediated human heart failure is associated with increased β1AR

It is not known whether anthracyclines which mediates deleterious cardiac remodeling affects cardiac βARs. Multiple studies including ours have shown that end-stage human heart failure dilated cardiomyopathy (DCM) patient samples are characterized by significant down-regulation of β1ARs^5, 6, 9^. Radio-ligand binding studies were performed to test whether βARs are altered in the end-stage human heart failure samples due to anthracycline (Adriamycin) cardiotoxicity or DCM patient samples [patient characteristics: **Table 1**]. Consistently, downregulation of β1ARs was observed in end-stage DCM samples^6, 9^ [**Fig. 1A**, grey bar, left panel] but surprisingly, there was significant increase in β1AR density in the human heart failure samples due to Adriamycin cardiotoxicity [**Fig.1A**, black bar, left panel]. Furthermore, β1AR density in the Adriamycin cardiotoxic human heart failure samples was significantly increased even compared to the non-failing controls [**Fig.1A** white vs. black bar, left panel]. However, β2AR density in the Adriamycin cardiotoxic heart failures samples was not appreciably different than non-failing controls [**Fig. 1A,** middle panel, white vs. black bar] which was significantly downregulated in end-stage DCM samples [**Fig.1A**, middle panel, grey bar]. While DCM heart failure samples showed significant downregulation of βARs [**Fig. 1A**, right panel, grey bar], the Adriamycin cardiotoxic heart failure samples showed minimal loss in βAR density compared to non-failing samples [**Fig.1A,** right panel, black vs. white bars]. These observations show that the chemotherapeutic agents may induce cardiac dilation/heart failure in otherwise healthy hearts by upregulating β1ARs, in contrast to heart failure etiology characterized by downregulation of β1ARs^5, 6, 9^. Consistently, studies have shown that β1AR signaling is pro-apoptotic^10^ and cardiomyocyte-specific overexpression of β1AR results in cardiac dilation and accelerated heart failure^5^.

**Figure:**
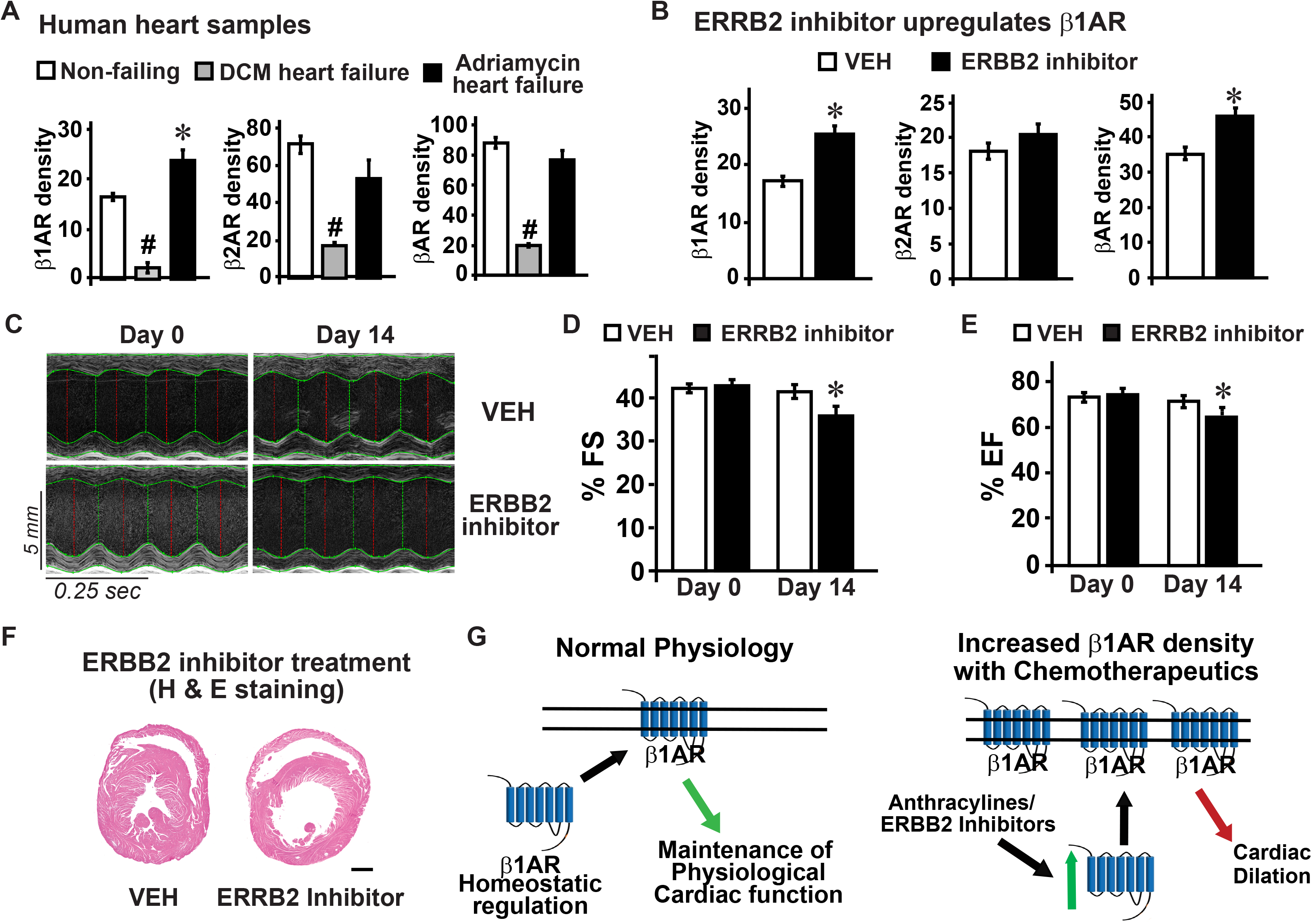
**A,** Radio-ligand binding in non-failing (NF) human heart (open bar) (n=5), dilated cardiomyopathy (DCM) (grey) (n=5) or Adriamycin-mediated heart failure (AHF) (black) (n=5). Left panel: β1AR density. #p<0.001 vs. NF and AHF; *p<0.001 vs. NF and DCM; Middle: β2AR density. #p<0.005 vs. NF and AHF; Right: Total βAR density. #p<0.005 vs. NF and AHF. **B,** β1AR density (left panel), β2AR density (middle) and total βAR density measured by radio-ligand binding on plasma membranes following Vehicle (VEH) or ERBB2 inhibitor (n=8), *p<0.0001 vs. Vehicle. Echocardiography for the VEH or ERBB2 inhibitor treated mice (n=11) (**C**) and associated functional assessment by % fractional shortening (%FS) (**D**) and % ejection fraction (EF) (**E**), *p<0.005 vs. other sample measures. **F,** Transverse heart section stained with H & E following VEH or ERBB2 inhibitor. Scale bar 1000 μm. **G,** Proposed schematic showing that anthracycline/ERBB2 inhibitor selectively elevates β1AR density and its signaling may underlie cardiotoxicity.

### ERBB2 inhibition results in increase of β1AR density

Since targeted chemotherapies like sunitinib, dasatinib or trastuzumab also leads to DCM, we tested whether alterations in the cardiac β1ARs could be a common underlying mechanism for cardiotoxicity. ERBB2 inhibition was performed by using AG 825 (a selective ERBB2 inhibitor) instead of trastuzumab/Herceptin as they are humanized antibodies which may initiate confounding immune response in mice. Radio-ligand binding studies showed modest yet, significant increase in β1AR density after ERBB2 inhibition in mice [**Fig. 1B**] with no appreciable changes in β2AR density [**Fig 1B**]. Consistent with the increased β1AR density, there was significant elevation in total βAR density [**Fig.1B**] following AG 825 administration. Interestingly, there was mild yet significant cardiac dysfunction following AG 825 treatment as observed by M-mode echocardiography and key measures of cardiac function % FS (fraction shortening) and % EF (ejection faction) [**Fig. 1 C, D** **&** **E**]. Furthermore, H & E staining reflected cardiac dilation following AG 825 treatment [**Fig. 1F**]. Similar to our previous studies^11^, appreciable cardiac dysfunction was not observed in females while males are susceptible to AG 825 within two weeks, an observation that is consistent with the role of estrogens in cardio-protection^12^. However, both males and females showed increase in β1AR density indicating that molecular events underlying the future deleterious outcomes precede the physiologic measures of cardiac function. Thus, increasing the timeline of AG 825 administration may lead to cardiac dysfunction in females. These findings indicate that chemotherapeutics (targeted/untargeted) may alter the homeostatic regulation of β1AR resulting in selective increase of β1AR density that promotes deleterious cardiac remodeling^5, 10^ [**Fig. 1G**]. These observations set the foundation to the conceptual idea that use of selective β1AR blockers may ameliorate cardiac dysfunction insulating the heart from chemotherapeutic cardiotoxicity.

## Abbreviations

EGFR: Epidermal growth factor receptor
ERBB2: Erythroblastic oncogene B
DCM: Dilated Cardiomyopathy
βAR: beta-adrenergic receptor

## Acknowledgements

The authors would like to thank Wendy Sweet and Dr. Christine S. Moravec for providing us end-stage DCM and anthracycline cardiotoxicity human heart failure samples. This work is in part, supported by Postdoctoral Fellowship Grant, AHA, 10POST3610049 (MKG).

